# Guard Cell-Enriched Phosphoproteome Reveals Phosphorylation of Endomembrane Proteins in Closed Stomata

**DOI:** 10.1101/2025.10.15.682613

**Authors:** Anne-Marie Pullen, Scott Lyons, Angie Mordant, Laura Herring, Belinda S. Akpa, Marcela Rojas-Pierce

## Abstract

Control of the stomatal aperture is multifaceted, involving a complex interplay of environmental cues and intracellular signaling pathways. It is well established that changes in ion gradients drive water movement into and out of the guard cell, thereby altering cell volume and modulating the opening or closing of the stomatal pore. These rapid responses are often regulated by phosphorylation cascades to efficiently transmit environmental status and either reduce water loss or enhance carbon assimilation. The role of endomembrane trafficking networks in stomatal dynamics is not well characterized. Here, we investigated the regulation of stomatal opening and closing by generating a proteome and phosphoproteome of guard cell-enriched tissue. This deep proteome captured a protein profile that was similar to previously characterized guard cell proteomes. The guard cell-enriched tissue with closed stomata showed greater levels of phosphorylation of proteins related to endomembrane trafficking and vacuoles when compared to both whole leaf tissue with closed stomata and guard cell-enriched tissue with open stomata. These results support the hypothesis that phosphorylation of endomembrane proteins may contribute to the regulation of stomatal movements.

## INTRODUCTION

The Arabidopsis stomatal complex consists of two guard cells surrounding a central pore. Guard cell volume directly influences pore size, with cell shrinking leading to stomatal closing and swelling leading to stomatal opening. The opening and closing of stomata are regulated by the movement of water between the apoplast and the guard cells. Water movement is largely driven by the passage of ions such as Ca^2+^, Cl^-^, H^+^, and K^+^ via ion transport channels within the plasma and vacuolar membranes (McAinsh et al. 1990; Andres et al. 2014; Hayashi et al. 2024; Yang et al. 2024).

Blue light is a key signal for stomatal opening, while abscisic acid (ABA) is a key signal for stomatal closing. The main pathway controlling blue light-induced stomatal opening involves the activity of PHOTOTROPIN (PHOT) 1 and 2, which are activated in response to blue light, leading to phosphorylation and activation of BLUE LIGHT SIGNALING1 (BLUS1) kinase (Takemiya et al. 2013; Liscum and Briggs 1995; Sullivan et al. 2008). BLUS1 activation ultimately results in activation of the H^+^ATPase pump, leading to a H^+^ proton gradient that results in water influx, guard cell swelling, and thus, stomatal opening (Schroeder et al. 1987). Some environmental stressors, such as water deficit, can trigger the accumulation of ABA, which relays stomatal closing signals through phosphorylation cascades (Schroeder et al. 2001; Kim et al. 2010). In the presence of ABA, the proteins PYRABACTIN RESISTANCE (PYR), PYRABACTIN-LIKE (PYL), and REGULATORY COMPONENT OF ABA RECEPTORS (RCAR) bind to a PROTEIN PHOSPHATASE 2C (PP2C) phosphatase, rendering it inactive (Park et al. 2009). This allows the OPEN STOMATA1 (OST1) kinase (Geiger et al. 2009; Pei et al. 1997) to activate the SLOW ANION CHANNEL ASSOCIATED 1 (SLAC1) S-type anion channel, resulting in K^+^ efflux (Pandey et al. 2007), a drop in guard cell volume, and stomatal closure (Mustilli et al. 2002).

Protein trafficking within the endomembrane system is regulated by suites of tethering complexes, soluble N-ethylmaleimide-sensitive factor attachment protein receptors (SNAREs), GTPases, and their effectors (Aniento et al. 2022; Qi et al. 2024; Rodriguez-Furlan et al. 2019). The vacuole is the largest plant cell organelle, and it is the destination for several trafficking pathways. There are two types of vacuoles in plants; protein storage vacuoles (PSV) are found in embryos, and lytic vacuoles are found in vegetative cells (Zheng and Staehelin 2011; Aniento et al. 2022). Trafficking to the vacuole can occur via the endocytic trafficking network or the anterograde vacuole pathway. The endocytic pathway removes proteins from the plasma membrane (PM) and delivers them to the *trans*-Golgi network (TGN), which in plants corresponds to the early endosome (Lam et al. 2007; Gonzalez Solis et al. 2022). The anterograde vacuole pathway traffics proteins from the endoplasmic reticulum (ER) to the vacuole (Herman and Schmidt 2004; Aniento et al. 2022). The TGN is an important protein sorting hub for proteins targeted for the PM or the vacuole (Kang et al. 2011; Griffiths and Simons 1986; Rosquete et al. 2018). Trafficking from the TGN to the vacuole requires maturation of TGN-derived multi-vesicular bodies (MVB) (Aniento et al. 2022), and MVB maturation is dependent on the sequential activity of RAB GTPases, specifically RAB5 and RAB7 (Cui et al. 2014; Ebine et al. 2014; Feng et al. 2017). Remodeling of the vacuole is mediated in part by endomembrane trafficking components and is a main contributor to changing guard cell volume and thus stomatal dynamics (Gao et al. 2005; Cao et al. 2022; Zheng et al. 2014; Mirasole et al. 2023). The vacuole within a guard cell is usually singular in open stomata and fragmented and convoluted in closed stomata (Gao et al. 2005; Cao et al. 2022; Zheng et al. 2014). Vacuole fusion is required for full stomatal pore opening, and this is regulated by vacuolar SNAREs and the HOMOTYPIC PROTEIN SORTING (HOPS) complex (Gao et al. 2005; Zheng et al. 2014; Takemoto et al. 2018; Brillada et al. 2018; Uemura et al. 2004; Uemura and Ueda 2014). The morphology of the vacuole is ultimately the result of vacuole fusion and MVB-vacuole trafficking (Cao et al. 2022; Brillada et al. 2018; Takemoto et al. 2018; Osorio-Navarro et al. 2025), and therefore, these pathways are important for understanding the role of vacuoles in stomatal movements.

Endocytic and exocytic trafficking pathways regulate the abundance of ion channels at the guard cell PM. The SNARE protein SYNTAXIN OF PLANTS 132 (SYP132) regulates the PM accumulation of H^+^-ATPase 1 (AHA1) in multiple cell types including guard cells (Xia et al. 2019). AHA1 localization is regulated in part by SYP132-mediated endocytosis when plants are under salinity stress (Baena et al. 2024), and interestingly, plants that overexpress SYP132 also have smaller stomatal apertures (Xia et al. 2019). Movement of K^+^ is at least partially regulated by the PM abundance of channels such as K^+^ TRANSPORTER OF ARABIDOPSIS THALIANA 1 (KAT1), and the trafficking of those channels is regulated by SYP121 (Leyman et al. 1999; Sutter et al. 2006; Sokolovski et al. 2008; Eisenach et al. 2012). Moreover, SYP121-mediated trafficking of KAT1 to the PM is regulated by phosphorylation in response to light (Ding et al. 2024). Overall, the endomembrane system is involved in the complex regulation of protein trafficking during stomatal opening and closing.

In this study, we explore the role of phosphorylation of the endomembrane system in open and closed stomata. We generated a guard cell-enriched proteome and phosphoproteome from *Arabidopsis thaliana* leaf epidermal fragments stimulated to have open or closed stomata. We found that guard cell-enriched tissue from closed stomata showed higher levels of phosphorylated proteins associated with trafficking and vacuole-related gene ontology (GO) terms compared to both, whole leaf tissue with closed stomata, or guard cell-enriched tissue with open stomata. This implies the existence of signaling pathways that result in phosphorylation of some trafficking proteins specifically in closed stomata.

## RESULTS AND DISCUSSION

### Generation of a guard cell-enriched proteome and phosphoproteome

A phosphoproteomic analysis of guard cell-enriched tissue was carried out to understand changes in phosphorylation of guard cells during stomatal opening and closing (Figure 1A). Mature Arabidopsis leaves were blended and epidermal fragments were filtered with a mesh to enrich for viable guard cells (Jalakas et al. 2017b). Angiosperm guard cells remain viable during this procedure due to their strong cell walls and lack of plasmodesmata (Bauer et al. 2013; Jalakas et al. 2017b; Shtein et al. 2017; Voss et al. 2018; Jalakas et al. 2017a; Sun et al. 2024; Geilfus et al. 2018; Rasouli et al. 2020). Epidermal fragments were exposed to conditions that induce stomatal opening or closing for 2 hours, and then immediately frozen in liquid nitrogen. A small aliquot was set aside at the end of the induction treatment, and stomatal opening or closing was confirmed via light microscopy (Figure 1A). Global proteomics and phosphoproteomics analysis was conducted on three technical replicates per condition using Tandem Mass Tag (TMT) labeling combined with high pH reversed-phase fractionation, FeNTA phosphopeptide enrichment and LC-MS/MS analysis. Phosphopeptide intensities were normalized to the abundance of the corresponding protein in each sample (*herein* adjusted value). Unless otherwise noted, all phosphopeptide data discussed below reflect these adjusted values. We previously reported differential phosphorylation of VPS39, a protein involved in vacuole fusion, between closed and open stomata in this dataset (Pullen et al. 2025). We also detected increased levels of phosphorylation of PHOT1 at S185 and S350, in the open samples and increased OST1 phosphorylation at S175 in the closed samples (Pullen et al. *Submitted*).

**Figure 1.**
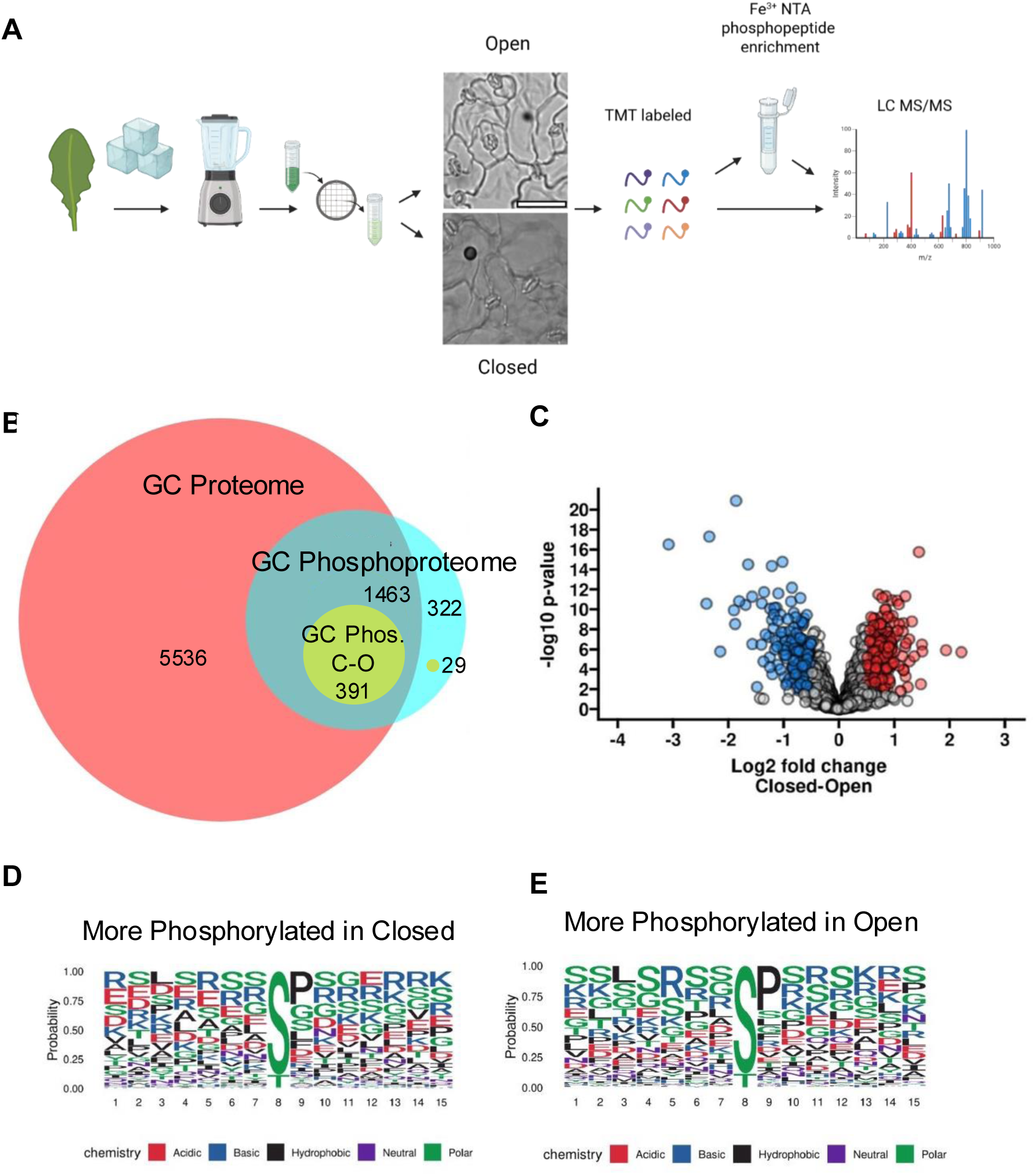
Overview of proteomics and phosphoproteomics study. A) Graphical protocol used for guard cell enriched phosphoproteomics. Scale bar at 64 µm. Drawing generated with BioRender. B) Comparison of proteome (GC Proteome, pink), phosphoproteome (GC Phosphoproteome, cyan), and differentially phosphorylated proteins between open and closed stomata (GC Phos. C-O, yellow). C) Volcano plot representing significantly differentially phosphorylated proteins from open and closed stomata. Proteins detected with higher (Red) or lower (blue) levels of phosphorylation in closed samples as represented as dots. D-E) Sequence logos show the over-represented motifs surrounding a centered PTM from peptides with higher levels of phosphorylation in closed samples (D, log2FC >0.5 and p-value <0.05), or higher levels of phosphorylation in open samples (E, log2FC <0.5 and p-value <0.05).

These modifications are consistent with prior reports (Sullivan et al. 2008; Wang et al. 2023), indicating that this is a valid dataset to identify changes in phosphorylation in open and closed stomata.

In total, 7,390 proteins were identified in the full proteome (Figure 1B, “GC proteome”, Table 1, Suppl File1) with only 17 instances of keratin contamination (not included in the 7,390). From the proteins identified, 143 proteins were more abundant in the closed samples as compared to the open, and 244 proteins were more abundant in the open samples as compared to the closed (Absolute Log2FC > 0.5, p-value < 0.05; Table 1). A total of 4,958 phosphorylated peptides were identified, mapping to 2,205 unique proteins across the full phosphoproteome dataset (Figure 1B, “GC Phosphoproteome”, Table 1, Suppl File2). Differential phosphoproteomics analysis revealed 420 unique proteins exhibiting significant changes (Absolute Log2 FC > 0.5 and p<0.05) between the closed and open samples, with a positive Log2FC indicating greater phosphorylation levels in the closed samples (Figure 1B, “GC Phos. C-O”, Figure 1C, Table 1, Suppl File2). Overall, 348 phosphopeptides were significantly increased in the closed condition and 222 phosphopeptides were significantly increased in the open condition (Table 1, Suppl File2). These results indicate a comprehensive profiling of the global proteome and phosphoproteome in guard cell–enriched tissue.

**Table 1.** Summary of proteome, and phosphoproteome. Phosphoproteome refers to identifiable proteins that are phosphorylated. Significance is determined as p< 0.05 and Log2FC < -0.5, or > 0.5.

|  | Unique proteins identified | Proteins more abundant in closed samples | Proteins more abundant in open samples | Total unique phosphorylated peptides | Peptides with increased phosphorylation in closed | Peptides with increased phosphorylation in open |
| --- | --- | --- | --- | --- | --- | --- |
| Proteome | 7390 | 143 | 244 |  |  |  |
| Phosphoproteome | 2205 |  |  | 4958 |  |  |
| Significant C-O Proteome | 420 |  |  | 570 | 348 | 222 |

### Motif analysis of phosphorylated proteins

Sequence alignment and motif enrichment analysis were used to identify phosphorylation motifs in the differentially phosphorylated protein dataset. Sequence logos were generated using the R program *[rmotifx](https://proteome-x.com/motif-x)* (Chou and Schwartz 2011) to identify over-represented motifs surrounding a centered residue that was significantly phosphorylated (Figure 1D, E). Comparisons between motifs found in the open or closed samples indicated that motifs including SPxR, SP, and RxxS were detected as enriched in the closed samples, with SPxR having the largest fold increase of 30.1, and SP and RxxS having fold increases of 6 and 5, respectively (Table 2). SP and RxxxxxxS motifs were enriched in the open samples with modest fold increases of 5.7 and 3.2, respectively. All of the previous sequences listed except for RxxxxxxS showed high motif scores in the 300s (Table 2). High motif scores indicate more statistically significant motif identification as well as motifs that are more specific to a particular kinase or kinase family. The minimal S/TP motif is the motif for Mitogen Activated Protein Kinases (MAPKs), some of which are implicated in ABA response (Amanchy et al. 2007; Umezawa et al. 2013). The RxxS motif corresponds to a more general motif in plants that is associated with SNF1-related protein kinase 2 (SnRK2s) such as OST1 (Furihata et al. 2006; Kobayashi et al. 2005; Umezawa et al. 2013) and calcium-dependent protein kinases (CDPKs) (Choi et al. 2005; Umezawa et al. 2013). Both SnRK2s and CDPKs are involved in guard cell signaling through phosphorylation (Mustilli et al. 2002; Hubbard et al. 2012; Geiger et al. 2009; Liu et al. 2022). The SP and RxxS motifs were enriched in ABA-treated leaves in a previous study (Umezawa et al. 2013). As shown in the motif analysis, many distinct phosphorylation motifs were over-represented in both the open and closed samples, reflecting the activity of diverse kinases involved in regulating stomatal opening and closing.

**Table 2.**
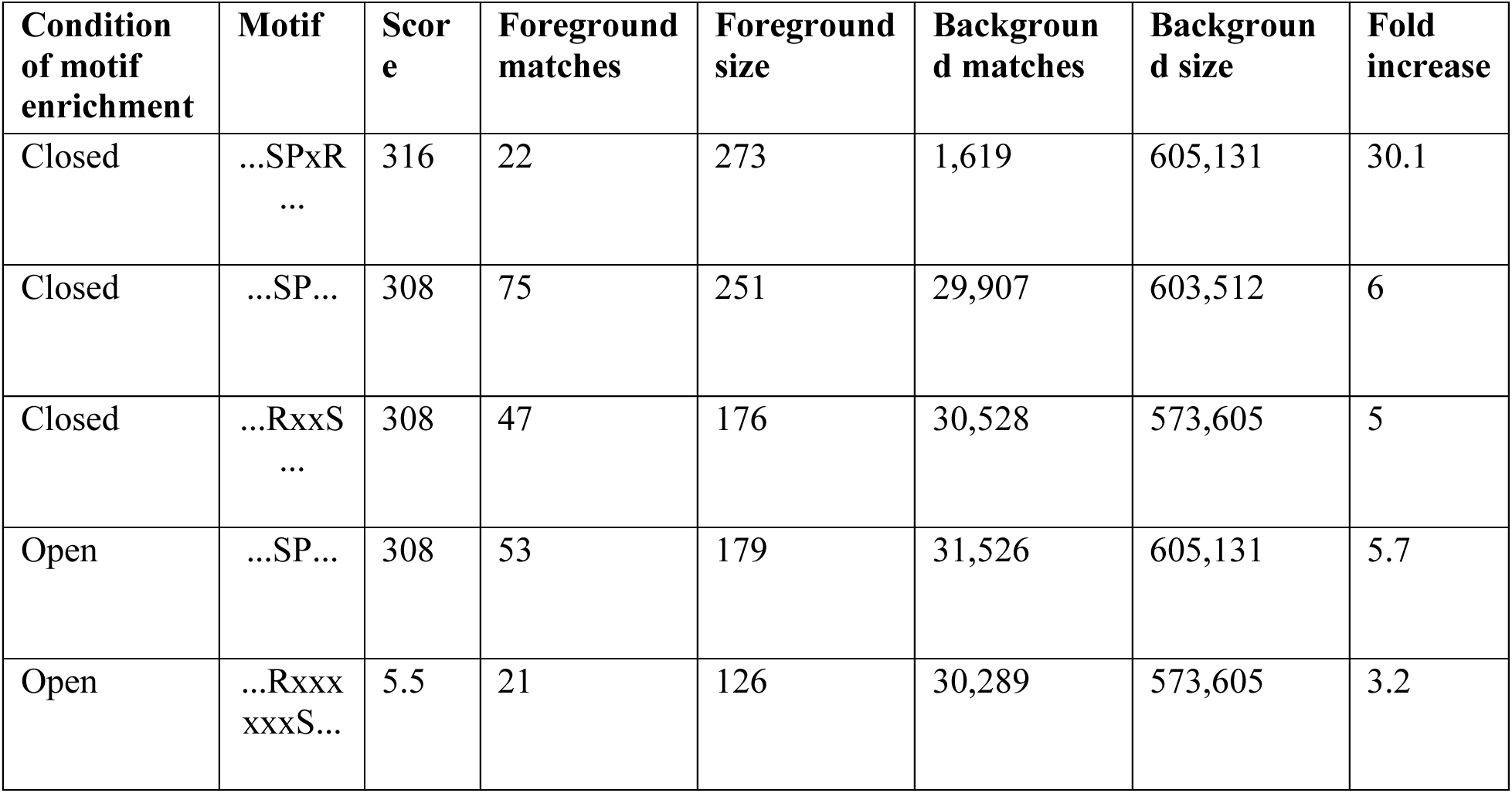
Top 5 phosphorylation motifs in guard-cell-enriched tissue. Table shows the top 5 motifs by score centered at S residue for the Closed-Open or Open-Closed comparison (Figure 1D, E). The first three rows are motifs that are overrepresented in the closed samples. The last 2 rows are motifs that are overrepresented in the open samples. The Score column refers to the motif score; the higher scores correspond to more statistically significant as well as more specific motifs. Matches refer to frequency of sequences matching this motif to either the foreground or background. Size refers to the total number of sequences. Fold increase is an indicator of the enrichment level of the extracted motifs.

| Condition of motif enrichment | Motif | Score | Foreground matches | Foreground size | Background matches | Background size | Fold increase |
| --- | --- | --- | --- | --- | --- | --- | --- |
| Closed | ...SPxR<br>... | 316 | 22 | 273 | 1,619 | 605,131 | 30.1 |
| Closed | ...SP... | 308 | 75 | 251 | 29,907 | 603,512 | 6 |
| Closed | ...RxxS<br>... | 308 | 47 | 176 | 30,528 | 573,605 | 5 |
| Open | ...SP... | 308 | 53 | 179 | 31,526 | 605,131 | 5.7 |
| Open | ...Rxxx<br>xxxS... | 5.5 | 21 | 126 | 30,289 | 573,605 | 3.2 |

### Comparison with previously published guard-cell proteomes

Previous studies investigated guard cell-enriched proteomes using epidermal fragments or protoplasts (David et al. 2020; Geilfus et al. 2018; Wang et al. 2023; Zhao et al. 2008). The proteins identified in our study overlap ∼80-87% with two smaller datasets (David et al. 2020; Geilfus et al. 2018) generated from similar epidermal fragment isolation methods (Figure 2, David and Geilfus, Table S1).

**Figure 2.**
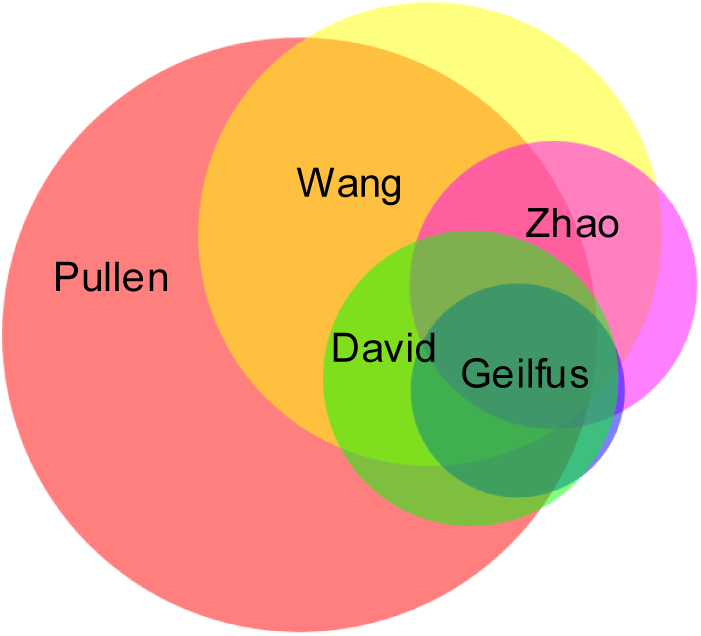
Proteome overlaps with previously described guard cell-specific proteomes. A) Guard-cell enriched proteome compared to two other studies with the same sample preparation of epidermal peels and crude protein extract (David et al. 2021; Geilfus et al., 2018). B) Guard-cell enriched proteome includes many proteins previously identified in guard cell protoplast or guard cell enriched proteomic studies. Wang et al., 2023; David et al., 2021; Geilfus et al., 2018; Zhao et al., 2008.

Our study also overlaps ∼75-80% with a smaller datasets derived from guard cell protoplasts (Figure 2, Wang and Zhao, Table S1) (Zhao et al. 2008; Wang et al. 2023). The high levels of overlap in protein identification indicate that the proteome profiles captured in our study are consistent with previously published studies. The greater depth of our proteome dataset provides new insights into guard cell-specific pathways that were not previously characterized. Notably, previous guard cell proteomics studies did not assess protein phosphorylation.

By using a TMT-based quantitative multiplexing method and avoiding protoplasting treatments, this study produced a deep proteome and phosphoproteome of intact and responsive guard cells. The larger proteomic coverage described in this study, compared to others with similar sample collection (Geilfus et al. 2018; David et al. 2020), may be due to the increased depth and coverage enabled by the TMT 10-plex strategy used for protein quantification (O’Connell et al. 2018; Pappireddi et al. 2019). This method uses an isobaric label that increases sample throughput. Additionally, the use of intact guard cells instead of protoplast may have improved the proteomic depth. Cell-protoplasting can result in lower depth of the proteome (Vu et al. 2024), upregulation of stress-related genes (Bates et al. 2012; Xu et al. 2021), and increased secretory activity to rebuild the cell wall (Denecke et al. 2012; Jeong et al. 2021; Bandmann et al. 2011). Finally, the phosphoproteome of protoplast samples could be altered by the additional time required for protoplasting (Rasouli et al. 2020).

One concern with using blended leaf fragments as a source of guard cell-enriched tissue is the possibility of detecting signal from other cell types. Some pavement and mesophyll cell fragments likely remained in the samples as observed by light microscopy (Figure 1A), which would result in a small percentage of identified proteins originating from mesophyll cells instead of guard cells. To test this, we compared our datasets to mesophyll or guard cell specific proteomes produced by Wang et al. (2023).

Wang et al. (2023) compared separate proteomes derived from guard cell or mesophyll cell protoplasts and characterized proteins specific to the guard cell proteome (Wang GCP) or mesophyll cell proteome (Wang MCP). Our proteome overlaps with 62.99% of the Wang GCP and also with 63.32% of the smaller Wang MCP (Figure 3A, Table S1). However, the overlap with mesophyll-specific proteins identified in Wang MCP represents only 1.96% of our full proteome. This result indicates that most proteins in our data are likely present in both mesophyll and guard cells. We hypothesize that differentially phosphorylated proteins in our dataset are more likely to be specific to the guard cells than any residual mesophyll cell proteins because of the responsiveness of guard cells to the opening and closing signals. From the differentially phosphorylated proteins from our dataset (Pullen Phos. C-O), thirty-one overlap with the Wang GCP, and only four overlap with the Wang MCP (Figure 3B). In a Fisher’s exact test, the proportion of either the MCP or GCP covered by the proteome is not significantly different (two tailed Fisher’s exact test p-value =1.00). However, the proportion of GCP proteins in the differentially regulated phosphoproteome is significantly greater than MCP proteins (two tailed Fisher’s exact test p= 0.0326, odds ratio MCP/GCP=0.33). Therefore, while most of the differentially phosphorylated proteins are likely found in both mesophyll and guard cells, the association analysis demonstrates that guard cell-specific proteins, and potentially guard cell signaling pathways, are enriched in the differentially regulated GC phosphoproteome.

**Figure 3.**
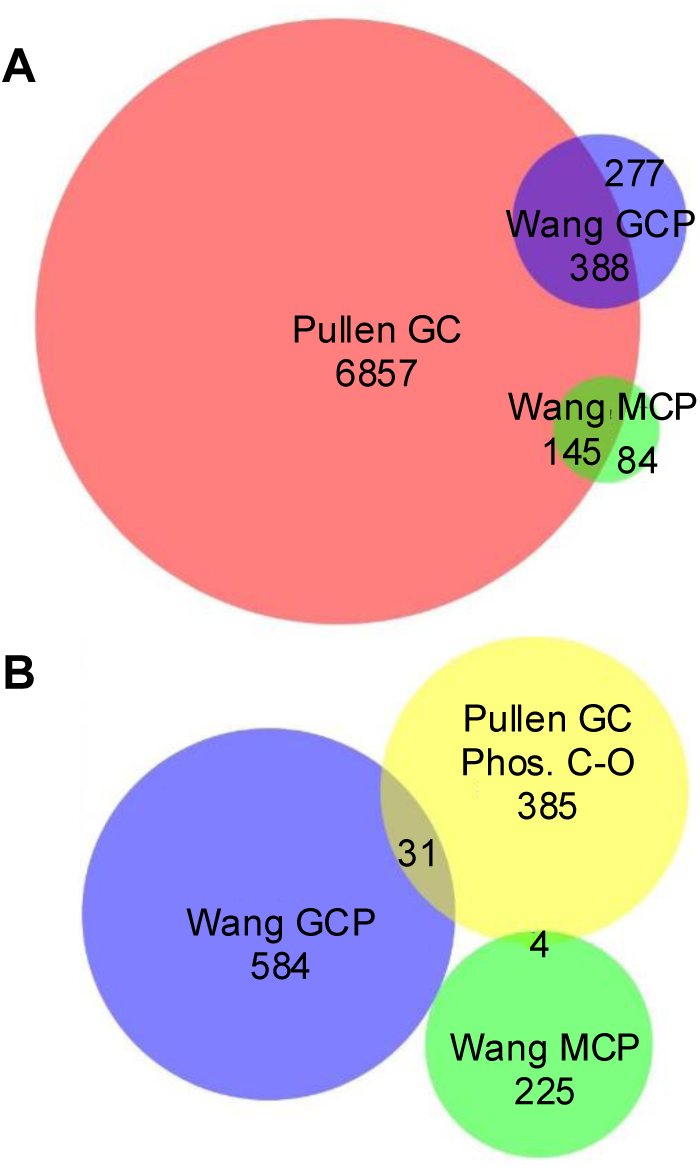
The differentially regulated phosphoproteome represents guard cell signaling. A) Comparison of the full GC proteome (pink) to guard cell (GCP) or mesophyll (MCP) specific proteins identified in Wang et al., 2023. B) Differentially phosphorylated proteins in this study (GC Phos. C-O, yellow) overlap with 31 guard cell-specific proteins (Wang GCP) and 4 mesophyll-specific proteins (Wang MCP).

### Functional enrichment analysis of guard cell protein phosphorylation reflects characterized stomatal pathways

To investigate the biological significance of our findings, we performed Gene Ontology (GO) analysis to identify overrepresented functional terms and pathways within our phospho-proteomics dataset. This analysis was conducted using the Singular Enrichment Analysis (SEA) tool from the Plant

Gene Set Annotation Database ([PlantGSAD](https://systemsbiology.cau.edu.cn/PlantGSEAv2/)) (Ma et al. 2022). Two separate SEA analyses were conducted on the differential phosphoprotein datasets for open and closed stomata. The top ten most enriched GO terms correspond to well-characterized pathways involved in stomatal movement (Figure 4). Four of the top ten terms are shared among the open and closed samples and correspond to broad terms including Cellular Process (GO:0009987), Cellular Metabolic Process (GO:0044237), Response to Stimulus (GO:0050896) and Response to Abiotic Stimulus (GO:0009628) (Figure 4).

**Figure 4.**
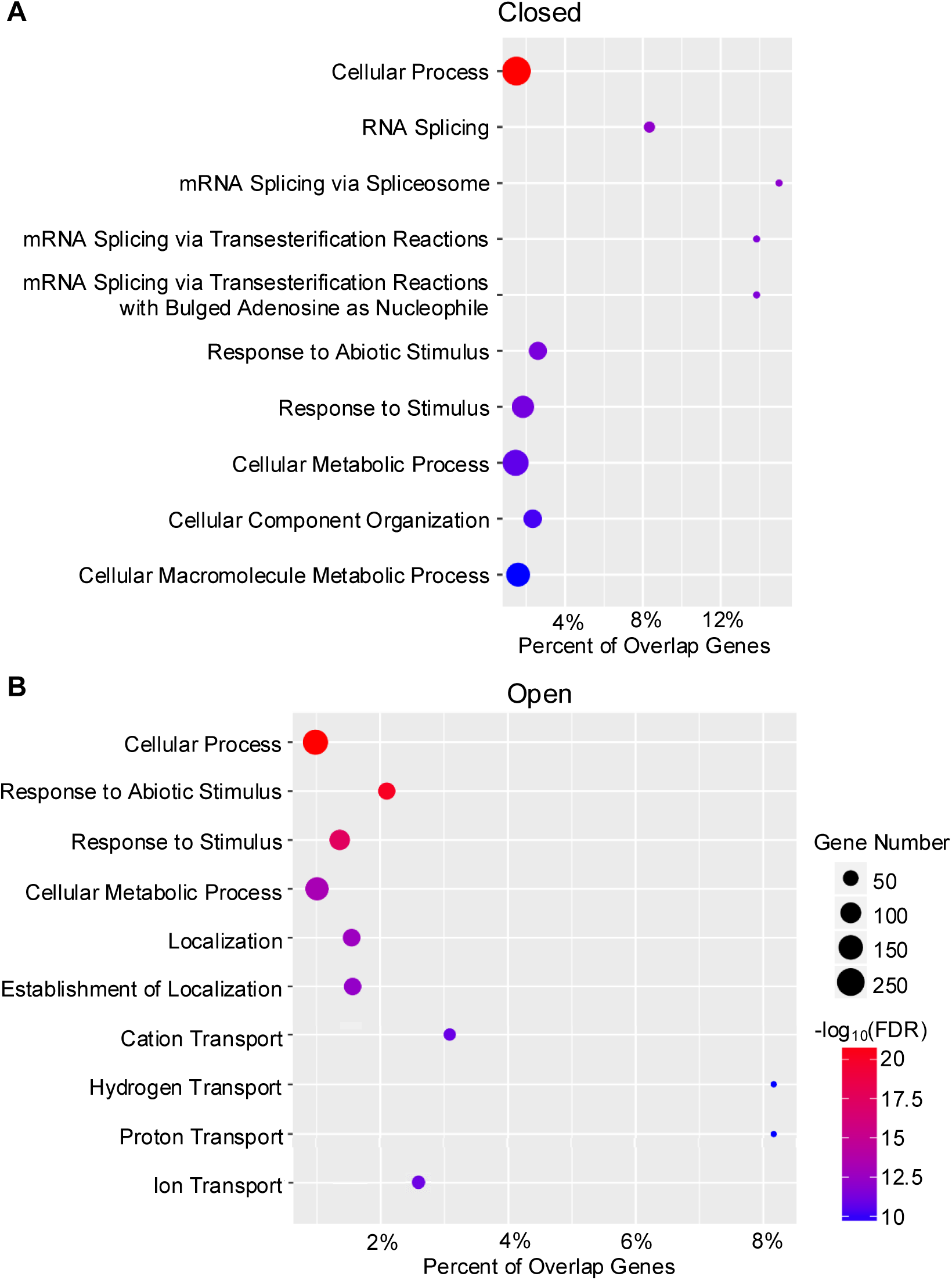
Top 10 Biological Process GO terms sorted by False Discovery Rate (FDR) for differentially phosphorylated proteins. (A) GO terms enriched for proteins of the peptides identified as more phosphorylated in the closed stomata samples (log2FC > 0.5, p< 0.05). (B) GO terms enriched for proteins of the peptides identified as more phosphorylated in the open stomata samples (log2FC < -0.5, p< 0.05).

Among the proteins with increased phosphorylation in the closed stomata samples, GO terms related to alternative splicing were overrepresented (Figure 4A). Alternative splicing is a well-documented response to environmental stress (Christensen et al. 1992; Larkin and Park 1999; Filichkin et al. 2010; Reddy et al. 2013; Marquez et al. 2012), and plant mRNA splicing machinery is also regulated by phosphorylation (de la Fuente van Bentem et al. 2006). Additionally, mimicking ABA responses by treating with pladienolide B (PB) leads to changes in alternative splicing and closure of stomata in fava bean and Arabidopsis (Ling et al. 2017). Therefore, enrichment for alternative splicing terms in the closed stomata phosphoproteome is consistent with previous reports. It is worth noting that these alternative splicing terms are still significant in the open samples, but with lower confidence as compared to the closed samples (Table S2).

Terms related to ion transport as well as enrichment for terms such as Localization (GO:0051179) or Establishment of Localization (GO:0051234) are overrepresented in the proteins with increased phosphorylation in the open stomata samples compared to the closed (Figure 4, Table S2). The presence of terms related to ion transport are consistent with the known phosphorylation events associated with ion transport activity in stomatal opening (Andres et al. 2014; Hayashi et al. 2024; Yang et al. 2024; Park et al. 2009). The terms related to localization, which refer to movement, tethering, and selective degradation, are equally enriched in both datasets (Figure 4, Table S2). The localization terms may hint at phosphorylation of trafficking proteins in stomatal opening and closing, but these terms are still very broad and are parent terms for the ion transport terms listed above (Binns et al. 2009). Analyzing the most overrepresented GO terms for the proteins with increased phosphorylation in the open or closed stomata samples provides a broad view of this dataset and largely confirms previously reported results.

### GO enrichment analysis reveals phosphorylation of trafficking proteins in closed stomata

To understand if the broad localization terms listed in Figure 4 are representative of more specific endomembrane trafficking terms present in the datasets, we conducted a SEA Compare analysis. We also wanted to determine whether terms related to endomembrane trafficking were enriched in the guard cell-enriched samples compared to other phosphoproteomes derived from whole leaf tissue. SEA Compare compiles the annotated GO terms for multiple SEA analysis and color codes the significance based on FDR (Ma et al. 2022). Using this tool, we compared the Pullen Phosphoproteome to proteins showing increased phosphorylation in Arabidopsis whole rosettes collected in the dark, one hour before (Giese Rosette Dark) or one hour after light exposure (Giese Rosette Light) (Giese et al. 2023). These two treatments replicate conditions that induce closing or opening of stomata, respectively.

The GO terms were searched for terms associated with stomatal closing (e.g. “ABA”) or stomatal opening (e.g. “blue light”) (Figure 5). The proteins with increased phosphorylation in the closed stomata samples (Pullen GC Closed) showed similar levels of enrichment for Stomatal Movement (GO:0010118) and slightly less enrichment for Response to ABA Stimulus (GO:0009737) when compared to the whole leaf tissue (Giese Rosette Dark) (Giese et al. 2023) (Figure 5). The proteins with increased phosphorylation in the open stomata samples (Pullen GC Open) showed enrichment for terms related to stomatal movement and blue light signaling, and Giese Rosette Light also shows enrichment for blue light signaling terms (Figure 5). These results indicate that both the Pullen GC and Giese phosphoproteomes represent phosphorylated protein populations that align to canonical terms associated with stomatal opening and closing (Takemiya et al. 2013; Park et al. 2009; Pandey et al. 2007; Giese et al. 2023).

**Figure 5.**
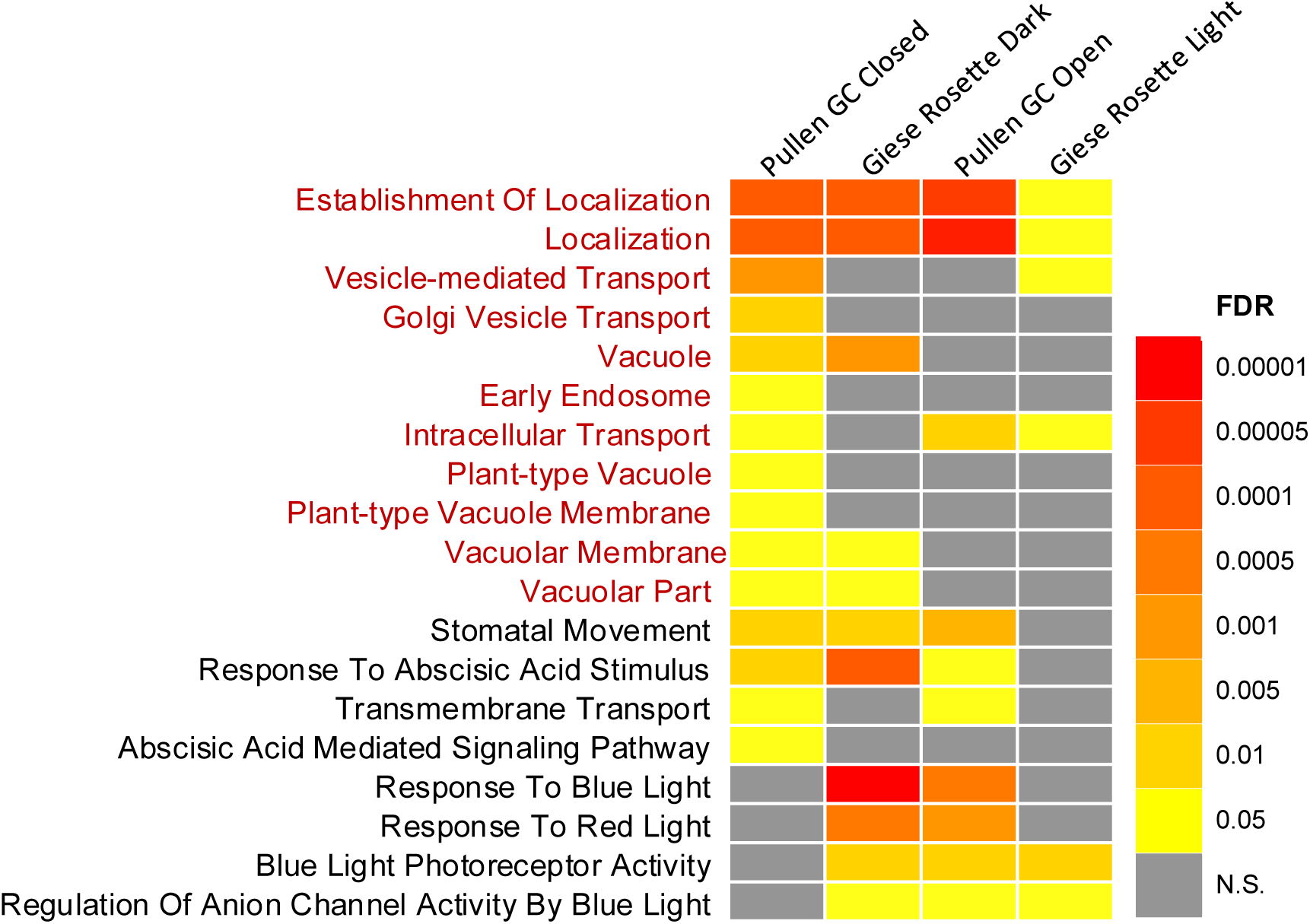
GO enriched terms of interest show closed guard cell specific phosphorylation of endomembrane trafficking and vacuole proteins. All comparisons are between peptides found to be significantly more phosphorylated in the closed stomata (Pullen GC Closed vs Giese Rosette Dark) or the open stomata state (Pullen GC Open vs Giese Rosette Light). Red text indicates terms used in the network analysis (Figure 6). For determining significance in phosphoproteome data, Log2FC >0.05 and p-value < 0.05 was used. FDR <0.05 was used for determining significance in SEA. Sources: Giese et al., 2023. SEA Compare conducted using PlantGSAD.

GO terms were searched for endomembrane system-related keywords (Table S3) including those containing the words vacuole, Golgi, and vesicle (Figure 5, Table S3). The broad terms Establishment of Localization and Localization were consistently highly enriched for all samples (Figure 5). Between the treatments, there is a trend wherein all samples exposed to closing conditions have increased enrichment for terms related to endomembrane trafficking and vacuoles when compared to samples exposed to opening conditions (Figure 5), with the closed guard cell-enriched tissue having the greatest enrichment in these terms. In increasing order of enrichment, Giese Rosette Light has the lowest enrichment for terms related to the vacuole and protein trafficking (Figure 5). The next highest level of enrichment is Pullen GC Open with 3 terms all at much higher confidence than Giese Rosette Light. Next, Giese Rosette Dark is enriched for 5 terms of interest. By comparison, the Pullen GC Closed data is enriched for 11 terms of interest with several terms only enriched in this dataset, specifically. These terms are Early Endosome (GO:0005769), Golgi-Vesicle Transport (GO:0048193), Plant-type Vacuole (GO:0000325) and Plant-type Vacuole Membrane (GO:0009705) (Figure 5). Terms related to protein trafficking and plant vacuoles are overrepresented in the closed stomata samples (Pullen GC Closed, Giese Rosette Dark, Figure 5). These terms are further enriched when analyzing proteins within guard cell tissue samples.

These data highlight a possible regulation of endomembrane trafficking by phosphorylation during stomatal movements, which could contribute to changes in stomatal dynamics.

### Network analysis of differentially phosphorylated proteins involved in the endomembrane system

We used the Search Tool for the Retrieval of Interacting Genes/Proteins ([STRING](https://www.string-db.org/)) database (Szklarczyk et al. 2023) to generate a putative interaction network of proteins with increased phosphorylation in the Pullen GC Closed sample that were also sorted into GO terms of interest (Table S3, Figure 6). The network used several interaction sources with a medium-high confidence level (0.6) for the minimum required interaction score; this cutoff balanced the number of possible interactions with the possibility of false positive interactions. These interactions are coded as experimentally determined protein-level interactions (experimentally determined), protein-level interactions listed in databases (curated database), documented co-expression for transcripts, genes, or proteins (conserved co-expression), protein homology, and protein names found in the same abstract (text-mining) (Figure 6) (Szklarczyk et al. 2023). A total of 73 proteins (nodes) were input into the database, and all non-connected nodes were removed resulting in a network of 16 nodes (Figure 6, Table S4). Among the proteins of interest in our network are KEEP ON GOING (KEG), RABA1g, BREFELDIN A (BFA)- INHIBITED GUANINE NUCLEOTIDE5 (BIG5), and GNOM-LIKE 1 (GNL1) (Figure 6). These proteins were selected because they are examples of vesicle trafficking proteins. Their possible connections to stomatal regulation are discussed below.

**Figure 6.**
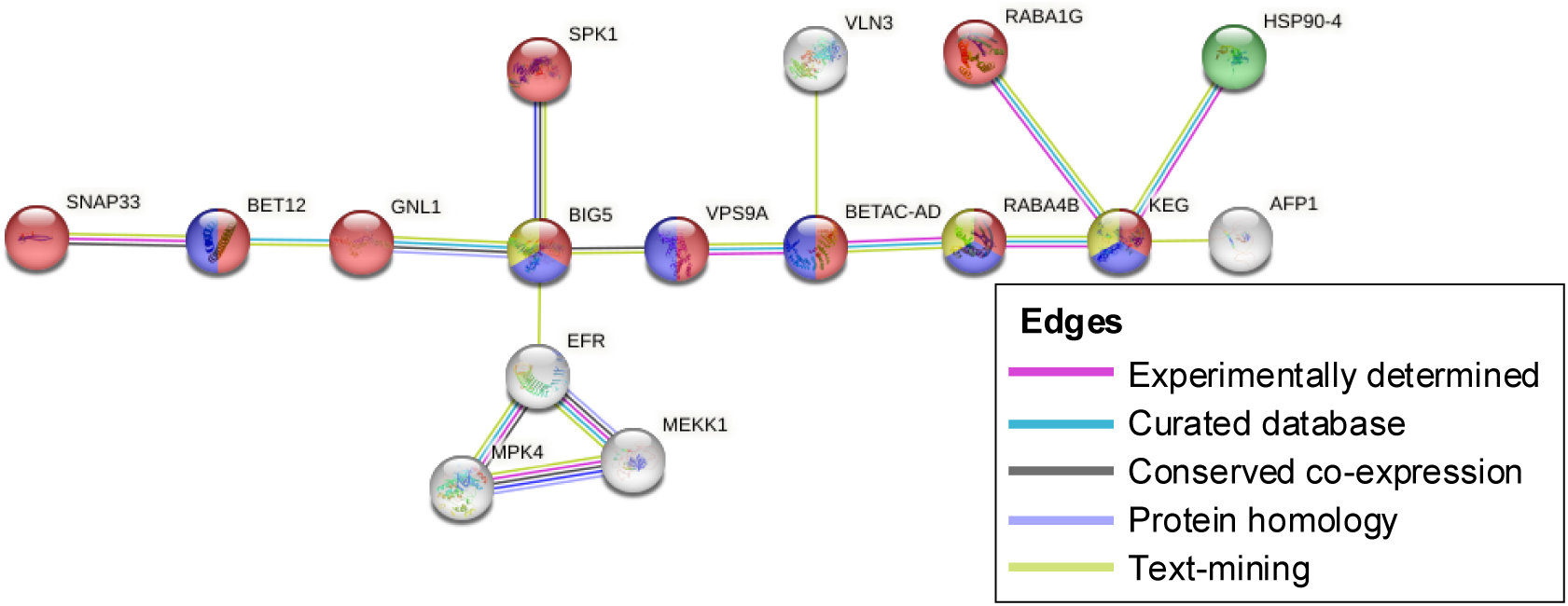
Interaction network of proteins that were enriched for phosphorylation in closed stomata. Diagram represents the Interaction Network of phosphorylated proteins found in terms of interest that were enriched in Pullen GC Closed data. Unconnected nodes were omitted. Number of edges: 16. Nodes were color coded based on GO terms such as Vesicle Mediated Transport (red), Intracellular transport (blue), Plant-type Vacuole (green), Early Endosome (yellow). Though the GO terms are the same between STRINGDB and Plant GSAD, some of the listings for the proteins differ. All proteins listed here were sorted into one of the 11 GO terms highlighted in figure 5. STRINGDB v 12. minimum required interaction score: 0.6. permalink: https://version-12-0.string-db.org/cgi/network?networkId=blpLS1HWn0aO

KEG is an E3 ubiquitin ligase that contains a kinase domain, localizes to the TGN, and regulates ABA signaling (Liu and Stone 2010; Stone et al. 2006; Gu and Innes 2011). In the presence of ABA, KEG is auto-ubiquitinated after being activated via phosphorylation (Liu and Stone 2010). The auto- ubiquitination of KEG is exploited by pathogens to disrupt secretory defense systems (Gu and Innes 2012; Liu and Stone 2010). In our dataset, KEG had increased phosphorylation levels in closed samples at S974 with a Log2FC of 0.55 (Table S4). KEG was also found to be phosphorylated by a negative regulator of plant immunity ENHANCED DISEASERESISTANCE1 (EDR1) at S974 in 4-week-old Arabidopsis leaf tissue (Gao et al. 2021), and this residue may be a target for KEG auto-ubiquitination and degradation (Gao et al. 2021). KEG also mediates the degradation of MITOGEN-ACTIVATED PROTEIN KINASE KINASE 4 (MKK4) and MKK5; two MAP kinase cascade members. These MKKs are involved in plant immunity and are active in guard cells by regulating stomatal patterning through phosphorylation of SPEECHLESS (SPCH) (Gao et al. 2021; Zhang et al. 2018). Whether the phosphorylation of KEG in our closed stomata dataset is related to its role in secretion, its interaction with MAP kinases or both, is yet to be determined.

RABA1g is a RAB GTPase involved in post-TGN trafficking (Stefano et al. 2015; Asaoka et al. 2013). RABA1g was found to interact with MITOGEN ACTIVATED PROTEIN KINASE 4 (MPK4) in a guard cell-enriched pull-down assay under stomatal opening conditions (Lin et al. 2023). MPK4 is highly expressed in guard cells and is thought to be involved in immune response because overexpression of MPK4 confers increased resistance to *Pseudomonas syringae* (Lin et al. 2023). RABA1g was detected once in our dataset with a Log2FC of 0.65 at S197 (Table S4). However, our putative interaction network does not show a direct interaction with MPK4 (Figure 6). MPK4 was also phosphorylated in our dataset, and this may indicate a kinase and RAB interaction in stomatal dynamics.

BIG5 is an ADP-ribosylation factor Guanine Nucleotide Exchange Factor (ARF-GEF) (Singh and Jurgens 2018) that regulates trafficking of PM proteins, including those involved in plant immunity such as BRASSINOSTERIOD SENSITIVE 1 (BRI1) (Xue et al. 2019). BIG5 localizes to the TGN (Nomura et al. 2011), and loss of function of BIG5 leads to greater susceptibility to *Pseudomonas syringae* (Nomura et al. 2011), possibly implicating an inability of stomata to close in response to pathogens. Interestingly, BIG5 had reduced levels of phosphorylation in 4-week-old leaves treated with salicylic acid and N- hydroxypipecolic acid, which typically causes stomatal closure (Zhang et al. 2024). BIG5 phosphorylation was detected three times in our dataset at S1435 (Log2FC 0.65), S1439 (Log2FC 0.53), and S1397 (Log2FC 0.59) (Table S4). None of these phosphorylation sites were detected by Zhang et al. (2024). BIG5 has multiple connections with other proteins in the network. This includes sharing edges with GNL1 (discussed below) and VACUOLAR PROTEIN SORTING9A (VPS9A), a GEF that can activate RAB5 family members (Goh et al. 2007). Given the role of BIG5 phosphorylation in responses to salicylic acid and BRI1 trafficking during stomatal closing (Shang et al. 2016), these novel residues could represent sites for control of trafficking by BIG5 in closed stomata.

GNL1 is an ARF-GEF involved in retrograde Golgi-to-ER trafficking (Richter et al. 2007; Jonsson et al. 2017) as well as trafficking of the protease Aleurain to the vacuole (Teh and Moore 2007). Traffic of the auxin efflux carrier PIN-FORMED 1 (PIN1) is at least partially regulated by GNL1 along with RABA1b and BIG5 (Tanaka et al. 2009; Teh and Moore 2007; Tanaka et al. 2013; Feraru et al. 2012). GNL1 was phosphorylated in our dataset once at S22 with a Log2FC of 0.66 (Table S4). In the network, GNL1 is connected to BIG5 and BET12, a Bet1/Sft1-like Qc SNARE that regulates early secretory transport (Uemura et al. 2004; Chung et al. 2018) (Figure 6). Though GNL1 has no direct known activity in stomata, its connection to BIG5 in our network and in literature may indicate an emerging network of phosphorylated trafficking proteins.

The GO term enrichment analysis indicates that trafficking proteins are enriched for phosphorylation in the Pullen Closed GC samples as compared to the others (Figure 5). These proteins are primarily associated with the TGN, which is a trafficking hub (Stone et al. 2006; Asaoka et al. 2013; Teh and Moore 2007; Nomura et al. 2011), and most have roles in secretory activities (Gu and Innes 2012; Liu and Stone 2010; Xue et al. 2019; Feraru et al. 2012) . Additionally, some of these proteins, like RABA1g, have documented functions in guard cells (Lin et al. 2023). The presence of these proteins in our network may indicate a putative role of phosphorylation of trafficking proteins in regulating closed stomata.

## CONCLUSIONS

In this paper, we describe a proteome and phosphoproteome of guard cell-enriched samples from open and closed stomata. The proteome demonstrated high levels of overlap with other proteomes derived from epidermal fragments (David et al. 2020; Geilfus et al. 2018) as well as guard cell protoplast (Wang et al. 2023; Zhao et al. 2008), yet many proteins were not previously found in other guard cell proteomes (Figure 2). We provide a deep phosphoproteome from guard cell-enriched tissue that can be a resource for further characterization of phosphorylation cascades in stomata.

Additionally, we found that guard cells have more phosphorylated proteins that were associated with GO terms related to trafficking and vacuoles in closed stomata, as compared to whole leaf tissue or guard cell-enriched tissue in open stomata. Therefore, signaling pathways that result in phosphorylation of some trafficking proteins specifically in closed guard cells are likely to exist.

## METHODS

### Plant materials and growth conditions

Arabidopsis ecotype Col-0 was used for all experiments. Arabidopsis seedlings were grown on 0.5 x Arabidopsis Growth media (AGM) comprised of 0.2% w/v Murashige and Skoog medium with MES (RPI M70300-50.0), 1% w/v Sucrose, pH to 5.7 with KOH, and 0.125% w/v GelRite (Research Products International G35020-250.0). Plates were incubated vertically for 5 days in a Percival CU35L5 growth chamber at a constant 22 °C with 120 µ mol m^-2^ s^-1^ of light intensity on a 16 h/8 h day/night cycle then transferred to pots containing MetroMix830 (Sun Gro Horticulture). The potted plants were grown on shelves in the lab space under LED lights (Monios-L T8 LED Grow Light) at 130 µmol m^-2^ s^-1^ of light intensity at soil surface on a 16 h/8 h day/night cycle and ambient temperature ranging from 21.1 °C to 23.5 °C with an average of 22 °C.

### Guard cell enrichment

Guard cell enrichment was conducted as outlined in Jalakas et al. (2017). A total of 144 WT Col-0 *Arabidopsis* plants were grown in individual pots to 4-weeks-old for the phosphoproteomic study. Plants were grown over 3 separate planting periods. Rosette leaves were excised, their mid-ribs were removed with scissors and fragments were kept in sterile ice-cold ddH_2_O water until processing. The leaf fragments were then blended with 4 °C sterile ddH_2_O water and ice. Samples were strained through 100 µm nylon mesh (Elko Filtering Co 06-210/33) to collect the epidermal fragments containing intact guard cells. For phosphoproteomics, the intact guard cells were then equilibrated with stomata buffer in the dark for 30 minutes, rinsed with sterile ddH_2_O water, and then incubated at 22 °C for 2 hours with opening buffer (10 mM MES pH 6.1, 50 mM KCl) (Schroeder et al. 1993) in the light or closing buffer (10 mM MES pH 6.1, 40 mM malate, 5 mM CaCl_2_) (Schroeder et al. 1993) supplemented with 50 µM ABA (Sigma Aldrich A1049) in the dark. To confirm stomatal opening or closing, a small aliquot of each extraction was mounted on slides and imaged using a compound microscope (Leica DM5000B and DMC4500 camera) using bright field with a 20x water objective. Closed samples were kept in dark and very quickly imaged at low light. Samples were then washed in sterile ddH_2_O water and flash frozen with liquid N for protein extraction (Jalakas et al. 2017b). This was repeated over 3 separate days with 48 plants used each day, and each enrichment was processed completely within one day.

### Protein sample preparation

Samples were ground in mortar and pestle with liquid nitrogen and protein was extracted in a lysis buffer consisting of 8M urea (Thermo Fisher 29700) in 50 mM Tris-HCl (VWR 97061-794) pH 7.8, 1x cOmplete™ EDTA-free Protease Inhibitor Cocktail (Sigma-Aldrich 11836170001) and 1x Pierce™ Phosphatase Inhibitor Mini Tablets cocktail (Thermo Fisher A32957). Samples were incubated on ice for 10 minutes, gently vortexed, and centrifuged at 8,000 × *g* at 4 °C for 10 minutes. The supernatant was further cleared by centrifugation at 12,000 × *g* at 4 °C for 5 minutes. The final supernatant for each condition was mixed to maximize total protein concentration and evenly divided between three technical replicates per condition (open or closed) and flash frozen. To measure total protein concentration, a small amount of each sample was aliquoted and used in a Bradford assay (AlfaAesar VWR AAJ61522-AP). Protein extraction from the frozen samples was conducted for all samples together on one day.

Protein analysis was carried out by the UNC Michael Hooker Metabolomics and Proteomics Core. Each sample was acetone precipitated, then pellets were reconstituted in 1M Urea and processed. Samples were then reduced with 5mM Dithiothreitol (Thermo Fisher 20290) for 45 minutes at 37 °C, alkylated with 15 mM iodoacetamide (Thermo Fisher A39271) for 30 minutes in the dark at room temperature. Samples were then digested with LysC (Wako, VWR 125-02543 1:50 w/w) for 2 hours at 37°C, followed by overnight trypsin (Sequencing Grade, Promega V5111, 1:50 w/w) digestion at 37°C. The resulting peptide samples were acidified, and desalted using desalting spin columns (Thermo Fisher 89852), and the eluates were dried via vacuum centrifugation. Peptide concentration was determined using Pierce Quantitative Colorimetric Peptide Assay (Thermo Fisher 23275).

A total of six samples (3 open and 3 closed) plus four ‘pooled’ samples (created by combining equal volumes of each sample) were labeled with TMT10 reagents (Thermo Fisher 90406) at room temperature. Prior to quenching, the labeling efficiency was evaluated by LC-MS/MS analysis. After confirming >98% efficiency, samples were quenched with 50% hydroxylamine (Sigma-Aldrich 467804) to a final concentration of 0.4%. Labeled peptide samples were combined 1:1, cleaned using a peptide desalting spin column (Thermo Fisher 89852), and dried via vacuum centrifugation. The TMT-labeled sample was offline fractionated over a 90 minute run, into 96 fractions by high pH reverse-phase HPLC (Agilent 1260) using an Agilent Zorbax 300 Extend-C18 column (3.5-µm, 4.6 × 250 mm) with mobile phase A containing 4.5 mM ammonium formate (pH 10) (Sigma-Aldrich 70221) in 2% (vol/vol) LC-MS grade acetonitrile, and mobile phase B containing 4.5 mM ammonium formate (pH 10) in 90% (vol/vol) LC-MS grade acetonitrile (Sigma-Aldrich 271004-100), as previously described (Klomp et al. 2024; Mertins et al. 2018). The 96 resulting fractions were then concatenated in a non-continuous manner into 24 fractions and 5% of each were aliquoted, dried down via vacuum centrifugation and stored at -80 °C until further analysis. The remaining 95% of each fraction was further concatenated into three fractions and dried down via vacuum centrifugation. For each fraction, phosphopeptides were enriched using the High Select Fe-NTA kit (Thermo Fisher A32992) per manufacturer protocol, then the Fe-NTA eluates were dried down via vacuum centrifugation and stored at -80 °C until further analysis. For each proteome sample, 205 µg of tryptic peptides were aliquoted and dried down.

All samples (24 proteome fractions; 3 phosphoproteome fractions, analyzed in duplicate) were analyzed by LC/MS/MS using an Ultimate 3000-Exploris480 (Thermo Fisher). Samples were injected onto an Ion Opticks Aurora C18 column (75 μm id × 15 cm, 1.6 μm particle size) and separated over a 70 (proteome) or 100 (phosphoproteome) minute method. The gradient for separation consisted of 5–42% mobile phase B at a 250 nl/minute flow rate, where mobile phase A was 0.1% formic acid in water and mobile phase B consisted of 0.1% formic acid in 80% acetonitrile The Exploris480 was operated in turboTMT mode with a cycle time of 3 seconds. Resolution for the precursor scan (m/z 375–1400) was set to 60,000 with an automatic gain control (AGC) target set to standard and a maximum injection time set to auto. MS2 scans (30,000 resolution) consisted of higher collision dissociate (HCD) set to 38; AGC target set to 300%; maximum injection time set to auto; isolation window of 0.7 Da; fixed first mass of 110 m/z.

### Proteomic data analysis

Raw data was processed by the Proteomics core using [Proteome Discoverer](https://www.thermofisher.com/in/en/home/industrial/mass-spectrometry/liquid-chromatography-mass-spectrometry-lc-ms/lc-ms-software/multi-omics-data-analysis/proteome-discoverer-software.html?erpType=Global_E1) (V3.1, Thermo Fisher) set to ‘reporter ion MS2’ with ‘10pex TMT’. Peak lists were searched against the *A. thaliana* database from Uniprot consisting of ∼16,000 reviewed protein sequences, appended with a common contaminants database, using Sequest HT within Proteome Discoverer. Data were searched with up to two missed trypsin cleavage sites, fixed modifications: TMT6plex peptide N-terminus and Lys, carbamidomethylation Cys, dynamic modification: N-terminal protein acetyl, oxidation Met. For phosphoproteome data, additional dynamic modification: phosphorylation Ser, Thr, Tyr. Precursor mass tolerance of 10 ppm and fragment mass tolerance of 0.02 Da (MS2). Peptide false discovery rate was set to 1%. Reporter abundance was calculated based on intensity, and for MS2 data, co-isolation threshold was set to 30.

Data post-processing analysis was performed in R (v 4.4.1) using the [MSstatsTMT](https://www.bioconductor.org/packages/release/bioc/html/MSstatsTMT.html) and MSstatsPTM packages (Kohler et al. 2023; Huang et al. 2020). Global median normalization on the spectrum level data was performed. Spectrum level data was summarized to the protein level using Tukey’s median polish.

To fulfill the requirements for imputation, there must be at least one nonmissing TMT channel for the same peptide spectrum match (PSM) in the run, and there must also be at least one nonmissing PSM from the same protein in the same TMT channel in that run. Statistical modeling and inference were performed using a linear mixed-effect model. The Benjamini-Hochberg (BH) method was used to adjust the p-values for multiple comparisons. The phosphoproteome abundance data has been adjusted (protein-adjusted phosphoproteome) to account for changes at the protein level using the MSstatsPTM package. Proportional Venn diagrams were created using the [Deep Venn](https://www.deepvenn.com/) web tool from (Hulsen 2022).

### Motif analysis

Phospho-motif analysis was performed using *[rmotifx](https://proteome-x.com/motif-x)*, an R programming language implementation of the motif-x software (Chou and Schwartz 2011) . The "score" column refers to the motif score, higher scores typically correspond to more statistically significant as well as more specific motifs. Matches refer to the frequency of sequences matching this motif to either the foreground or background. Size refers to the total number of sequences. Fold increase is an indicator of the enrichment level of the extracted motifs.

### Gene ontology

[PlantGSAD](https://systemsbiology.cau.edu.cn/PlantGSEAv2/) was used to create gene ontology data including SEA, their dotplot drawer tool, and the SEA compare tool to compare GO term enrichment between studies (Ma et al. 2022). Terms of interest were selected to capture functions related to endomembrane trafficking and vacuoles. This included searching for terms containing: vacuole, inositide, inositol, vesicle, localization, trafficking, traffic, Golgi, endosome, clathrin, endo and exocytosis. Additional terms related to stomatal opening and closing were also searched including red, blue, light, stomata, ABA and abscisic acid.

### Network analysis

A list of proteins sorted into 11 GO terms of interest were compiled and duplicates shared between GO terms were removed (Table S3). The remaining list of 78 proteins was loaded into the [STRING](https://string-db.org/) database v12 (Szklarczyk et al. 2023). The categories of interaction used in this network as defined by STRING are as follows. “Known Interactions” are from curated databases, and “experimentally determined” interactions are results from published papers. The “Others” category includes co-expression, protein homology, and text-mining data. These interactions can include data from homologous proteins in other organisms like *Saccharomyces cerevisiae.* A minimum required interaction score of 0.6 was used.

Generally, interaction scores are divided into low confidence (0.15 or better) medium confidence (0.4 or better), high confidence (0.7 or better) and highest confidence (0.9 or better). The score of 0.6 was chosen to balance the ability to detect interactions with the reduction of false positives.

## ACKNOWLEDGEMENTS

This work was supported by the National Science Foundation (MCB-1918746 to M.R.P. and B.S.A.), by the Research Capacity Fund (HATCH), project award no. 7005574 (to M.R.P) from the U.S. Department of Agriculture’s National Institute of Food and Agriculture, and in part by NCSU through access to the Cell and Molecular Imaging Facility.

