## Supplementary material for "Guard Cell-Enriched Phosphoproteome Reveals Phosphorylation of Endomembrane Proteins in Closed Stomata": Suplemental Tables

### SUPPLEMENTAL TABLES

**Table S1. Comparisons with previous guard cell proteomic studies**. Wang MCP corresponds to the mesophyll specific proteins identified in Wang et al. 2023. GCP stands for the guard cell specific proteins identified in Wang et al., 2023.

| | **This Study** | **Zhao et al., 2008** | **David et al., 2021** | **Geilfus et al., 2018** | **Wang et al., 2023** | **Wang MCP** | **Wang GCP** |
|---|---|---|---|---|---|---|---|
| total protein ID per study | 7390 | 1732 | 1827 | 959 | 4492 | 229 | 616 |
| % overlap with this study | n/a | 80.1% | 80.7% | 87.2% | 74.4% | 63.3% | 62.9% |
| Sample Preparation | intact blended GC | GC protoplast | intact blended GC | intact GC peels | GC protoplast | MC protoplast | GC protoplast |

**Table S2. Comparison of top 10 GO terms.** Comparison of top 10 GO terms from Figure 4 sorted by FDR. Significance is defined as FDR <0.05.

| **Gene Ontology Term** | **Pullen GC Closed FDR** | **Pullen GC Open FDR** |
|---|---|---|
| Cellular Process | 1.16E-22 | 1.60E-13 |
| RNA Splicing | 2.08E-13 | 6.05E-05 |
| MRNA Splicing, Via Spliceosome | 3.02E-13 | 0.00102 |
| RNA Splicing, Via Transesterification Reactions | 9.79E-13 | 0.00153 |
| RNA Splicing, Via Transesterification Reactions With Bulged Adenosine As Nucleophile | 9.79E-13 | 0.00153 |
| Response To Abiotic Stimulus | 1.54E-12 | 4.52E-13 |
| Response To Stimulus | 2.37E-12 | 1.15E-11 |
| Cellular Metabolic Process | 8.09E-12 | 2.02E-09 |
| Cellular Component Organization | 2.45E-11 | 0.00021 |
| Cellular Macromolecule Metabolic Process | 7.96E-11 | 3.12E-06 |
| Establishment Of Localization | 2.51E-07 | 6.27E-09 |
| Localization | 2.68E-07 | 3.92E-09 |
| Cation Transport | 0.0156 | 3.39E-08 |
| Hydrogen Transport | 1 | 1.42E-07 |
| Proton Transport | 1 | 1.42E-07 |
| (Monoatomic) ION TRANSPORT | 0.00285 | 3.91E-07 |

**Table S3. GO terms used in SEA compare analysis and network.** GO terms from SEA compare analysis used in the network and Figure 5. Lists of proteins that sorted into these GO terms from the SEA Compare Analysis were downloaded. Duplicates between lists were removed and a list of 73 proteins was uploaded to STRING (Szklarczyk et al. 2023) for network analysis.

| **GO Term** | **GO number** |
|---|---|
| Establishment of Localization | GO:0051234 |
| Localization | GO:0051179 |
| Vesicle Mediated Transport | GO:0016192 |
| Golgi Vesicle Transport | GO:0048193 |
| Intracellular Transport | GO:0046907 |
| Early Endosome | GO:0005769 |
| Vacuole | GO:0005773 |
| Plant Type Vacuole | GO:0000325 |
| Plant Type Vacuole Membrane | GO:0009705 |
| Vacuolar Membrane | GO:0005774 |
| Vacuolar Part | GO:0044437 (obsolete) |

**Table S4. Phosphoproteomic data of network proteins**. A table of peptides belonging to the 16 proteins found in Figure 6.

| Gene Symbol | Protein Accession | Araport ID | PTM site | Log2FC Closed-Open | p-value Closed-Open |
|---|---|---|---|---|---|
| SNAP33 | Q9S7P9 | AT5G61210 | S194 | 0.72 | 2.58E-06 |
| BET12 | Q94CG2 | AT4G14455 | S30; S31 | 0.59 | 4.28E-02 |
| GNL1 | Q9FLY5 | AT5G39500 | S22 | 0.66 | 9.88E-03 |
| SPK1 | Q8SAB7 | AT4G16340 | S1051 | 0.51 | 3.49E-05 |
| BIG5 | F4IXW2 | AT3G43300 | S1435 | 0.65 | 3.93E-04 |
| BIG5 | F4IXW2 | AT3G43300 | S1439 | 0.53 | 6.59E-04 |
| BIG5 | F4IXW2 | AT3G43300 | S1397 | 0.59 | 1.92E-02 |
| EFR | C0LGT6 | AT5G20480 | S1010 | 0.53 | 7.96E-04 |
| MPK4; MPK11 | Q39024; Q9LMM5 | AT4G01370 | S195 | 0.99 | 3.62E-06 |
| MEKK1 | Q39008 | AT4G08500 | S111 | 0.58 | 1.11E-04 |
| VPS9A | Q9LT31 | AT3G19770 | S292 | 0.57 | 1.77E-04 |
| VLN3 | O81645 | AT3G57410 | S779 | 0.75 | 6.27E-05 |
| VLN3 | O81645 | AT3G57410 | S779; S781 | 0.67 | 7.58E-04 |
| BETAC-AD | O81742 | AT4G23460 | T660 | 0.57 | 1.86E-02 |
| RABA4B | Q9SMQ6 | AT4G39990 | S193 | 1.51 | 4.00E-05 |
| RABA4B | Q9SMQ6 | AT4G39990 | S193 | 1.36 | 9.29E-05 |
| RABA4B | Q9SMQ6 | AT4G39990 | S60 | 1.28 | 2.35E-04 |
| RABA1G | Q9LK99 | AT3G15060 | S197 | 0.65 | 2.08E-06 |
| KEG | Q9FY48 | AT5G13530 | S974 | 0.55 | 2.15E-05 |
| HSP90-4; HSP90-2; HSP90-3 | O03986; P55737; P51818 | AT5G56000 | S288 | 0.82 | 6.35E-06 |
| HSP90-4; HSP90-2; HSP90-3 | O03986; P55737; P51818 | AT5G56000 | S219 | 0.67 | 3.76E-03 |
| AFP1 | Q9LQ98 | AT1G69260 | S115 | 1.06 | 8.75E-07 |
